# 3 Directional Inception-ResUNet: deep spatial feature learning for multichannel singing voice separation with distortion

**DOI:** 10.1101/2023.07.20.549865

**Authors:** DaDong Wang, Jie Wang, MingChen Sun

## Abstract

Singing voice separation on robots faces the problem of interpreting ambiguous auditory signals. The acoustic signal, which the humanoid robot perceives through its onboard microphones, is a mixture of singing voice, music, and noise, with distortion, attenuation, and reverberation. In this paper, we used the 3 directional Inception-Resnet structure in the U-shaped encoding and decoding network to improve the utilization of the spatial and spectral information of the spectrograms.

Multi-objectives were used to train the model: magnitude consistency loss, phase consistency loss, and magnitude correlation consistency loss. We recorded the singing voice and accompaniment derived from the MIR-1k datasets with NAO robots and synthesized the 10-channel datasets for training the model. The experimental results show that the proposed model trained by multi-objective reaches an average NSDR of 11.55db on the test datasets, which outperforms the comparison model.

**Author summary:** The mixture in the real singing voice separation is always mixed with noise and distortion. In this paper, the acoustic signals with distortion and noise perceived by the robot are used to study the separation of singing voices in real scenes. This paper described how to synthesize the training datasets, proposed a 3 directional Inception-ResUNet structure for multichannel singing voice separation, and adopted multi-objectives including magnitude correlation consistency loss to train the model. The experimental results showed that the magnitude correlation consistency loss reduces distortions, the proposed model achieves better performance than the compared models.

## Introduction

Music training implemented by robots is more interesting than other devices. The robot has to complete accompaniment, intelligent music synthesis, interactive scoring, singing voice separation, lyrics synchronization, etc. The singing voice separation is the basis of other functions and is also important to improve the accuracy of robot speech recognition. Robots usually have 2-4 microphones. Recently, more and more researches focus on exploring the multi-microphone source separation in real-world applications [1]. Due to their physiological structure, humans can easily distinguish between singing voice and instrumental accompaniment when listening to a song. However, this is a challenge for machine learning models or deep learning models, since singing voice and accompaniment are strongly correlated in time and frequency. Moreover, multi-channel singing voice separation is more challenging due to model complexity, background noise, microphone distortion, and other factors.

There are two major approaches for multichannel source separation in the early stages: microphone array processing and blind source separation(BSS) [2]. The BSS approaches usually exploit the statistical characteristics of the mixture of the singing voice, accompaniment, and noise, while the microphone array processing approaches usually concern the signal models. The ideas between the two major approaches often borrow from each other. Recently, supervised singing voice separation using deep neural networks(DNN) has received widespread attention from researchers with great success [3]. Typically, these methods [4] [5] [6] [7]learn a mapping function from singing voice features to separation targets through supervised learning algorithms. Compared to signal models, deep learning can automatically extract the most powerful features of the singing voice in the mixture. The deep learning models can process the original high-dimensional data without knowledge requirements for feature design, mine the structured features in the singing voice, and output the structured prediction. Using DNNs in training had emerged as a promising trend in the field of microphone array processing and BSS [8] [9] [10] [21].

The researchers have proposed many deep learning models for BSS including recurrent neural networks(RNN) [4] [5] [11], convolutional neural networks(CNN) [12] [13], U-Net [6] [14] [15], long short-term memory(LSTM) [16] [17], generative adversarial networks (GAN) [18] [19] [20], etc. The results show that a DNN model trained with the singing voices of dozens of singers can separate the singing voices of others. Most DNN models [4]-[15] [17]-[20] deal with the time-frequency(T-F) domain spectrograms generated from the short-term Fourier transform(STFT). DNN models extract the spectral characteristics of the singing voice and accompaniment.

The microphone array processing approaches traditionally utilized spatial cues such as geometry-based information to construct the signal models [23] [24] [25]. In recent years, DNNs have emerged in multi-microphone array processing approaches for parameter estimation [26] [27] [28], spatial and spectral feature extraction [9] [29] [30]. Joint modeling of spatial and spectral information potentially improves the separation performance [2]. However, most previous DNN-based approaches do not fully utilize the spatial and spectral information of the spectrograms or lost part of the information in training, which leads to a certain residual noise in the separation results. On the other hand, selecting a proper training target for the singing voice separation is difficult. Single-objective, such as MSE or L1 loss, converges faster, but the results are not necessarily the best, and the problem with multi-objective training is how to balance multiple objectives.

Our contributions mainly include three aspects: (1)To improve the utilization of spatial and spectral information of the spectrograms, we proposed a 3-directional Inception-Resnet structure as the first layer of the U-shaped encoding and decoding network. (2)We trained the model with magnitude L1 Loss, correlation coefficient, and phase L1 Loss and obtained better separation performance. (3)We produced a kind of 10 channels datasets that can be used to test multi-channel singing voice separation algorithms.

### Related Work

DNN-based multi-channel singing voice separation is a supervised learning problem. This section introduces the system framework, training targets, and network models.

#### System framework

Most DNN-based singing voice separation consists of three stages [4]-[15] [17]-[20] [30] [31]:

1. Time-frequency transform. The time domain signals of the singing voice and mixture are decomposed into two-dimensional time-frequency-domain spectrograms by short-term Fourier transform(STFT), respectively.
2. Separation of singing voice using the DNN model. The model output is a soft mask that separates the mixture spectrogram into a singing voice and a non-voice spectrogram.
3. Frequency-time transform. The target singing voices in the time domain are reconstructed from the mixture spectrogram multiplied element-wise with the mask by inverse short-time Fourier transform (ISTFT).

#### Training targets

The STFT spectrograms expressed in complex numbers consist of two kinds of information: magnitude and phase. As the training data must be scalar, some studies use magnitudes, namely the modulus of complex numbers [6] [15] [20] [9]. The training target can be a spectral magnitude mask(SMM), defined as the magnitudes of the clean singing voice divided by that of mixture [32]. The magnitude is the square of the sum of squares between the real and imaginary components. Obviously, the phase cannot be calculated from the magnitude scalar. If the loss function does not include phases, SMM will lose the phase completely. The phase-sensitive mask(PSM) [22] and complex ideal ratio mask(cIRM) [33] contain the phase in the target. The PSM extends the SMM by multiplying *cosθ*, where *θ* denotes the difference between the clean singing voice and the mixture phase. The cIRM is defined as the spectrograms of the clean singing voice divided by that of the mixture. The PSM and cIRM potentially yield better separation performance than the SMM.

Compared to the monaural singing voice separation, the multichannel singing voice separation can use spatial in addition to spectral information. The interchannel level difference(ILD) and interchannel phase difference(IPD) can be used in training [34] [9] [35]. The mixture of the ILD, IPD, and spectrogram can still use SMM, PSM, or cIRM as training targets.

#### DNN models

The multichannel singing voice separation is a regression problem. Recently, most methods adopt an encoder-decoder structure to solve the problem. The encoder structure typically uses convolution and pooling to extract spectral features of the clean singing voice from the mixture of the ILD, IPD, and spectrogram, while the decoder structure uses deconvolution to recover the spectrogram of the clean singing voice. As the downsampling yields detail loss, usually, the upsampling is compensated with skip connections which connect the spectrogram with the result of upsampling in the same layer. Since Wang et al. used a 4-layer DNN to separate source [41], dozens of methods for singing voice separation using DNNs such as CNNs, RNNs, and various variants have been proposed [3]. After Jansson used U-Net for singing voice separation that surpassed the previous methods [6] [14], a few improved versions based on U-Net architecture achieved better performance [7] [9] [15] [46]. Our proposed model is also a variant of U-Net.

## Method

### Problem formulation

The time domain mixture signal gathered by the *i*th microphone can be defined by:

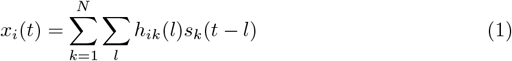

where *N* denotes the number of sources, *M* denotes the number of microphones, *k* ∈ 1, 2, …, *M k*≠ *i, s*_*k*_(*t*) denotes the signals recorded by the *i*th microphone from the *k*th source, *h*_*ik*_(*l*) denotes the impulse response from the *k*th source to the *i*th microphone, *l* denotes the impulse index. The spectrogram at time-frequency point (t,f) of *x*_*i*_(*t*) can be approximated as [40]:

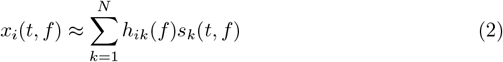

where *h*_*ik*_(*f*) is the frequency response from the *k*th source, *s*_*k*_(*t, f*) is the STFT of *s*_*k*_(*t*). As multiple noise sounds can be modeled as a single source [2], we denote with 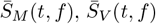 and 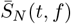 the spectrogram of music, singing voice and noise recorded by the *i*th microphone respectively. *x*_*i*_(*t, f*) can be described by:

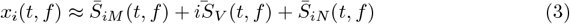

where 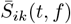 represents *h*_*ik*_(*f*)*s*_*k*_(*t, f*).

Taking the *i*th microphone as a reference, we used two relative transfer functions between the *i*th microphone and the *k*th microphone as [23].The *i*th microphone and the *k*th microphone interchannel level differences(ILD) is defined as:

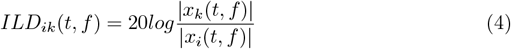

The *i*th microphone and the *k*th microphone phase differences(IPD) is calculated as:

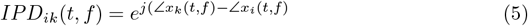

We concatenate the spatial cues *ILD*, the real components of *IPD*,the imaginary components of *IPD*, and the spectral features of each microphones {*ILD*_*ik*_(*t, f*), *IPD*_*ik*_(*t, f*).*real, IPD*_*ik*_(*t, f*).*imag*, 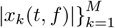 to form the input features.

The prediction targets 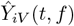 are the magnitude spectrograms of the singing voice. After training, the DNN model’s output predictions, which is a time-frequency mask, can predict the magnitude spectrograms of the target singing voice from the multichannel spectrograms. The mask is defined as [22]:

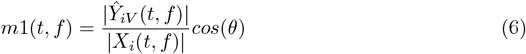

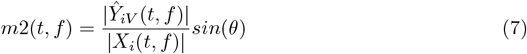

where *X*_*i*_ denotes the spectrograms of the reference microphone, *f* = 1, 2, 3, …, *F*, stands for different frequencies, and *θ* denotes the difference of the predicted singing voice phase and mixture phase of the reference microphone. After the soft mask is calculated, it is applied to *X*_*i*_ to estimate the predicted separation spectrograms 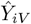.

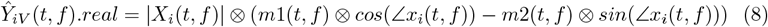

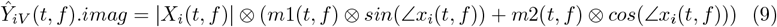

where ⊗ stands for element-wise operation.

### loss function

To avoid the phase independence of the predicted spectrograms, we used a kind of hybrid phase-dependent loss function to train *m*1(*t, f*) and *m*2(*t, f*).

(1) magnitude consistency loss *Loss*_*M*_

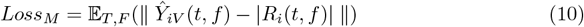

where *R*_*i*_ denotes the normalized spectrograms of the target clean singing voice, 𝔼_*T,F*_ is the mathematical expectation in data domains T and F.

(2) phase consistency loss *Loss*_*P*_

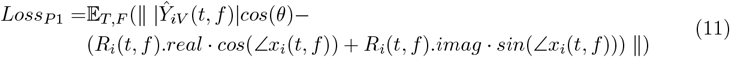

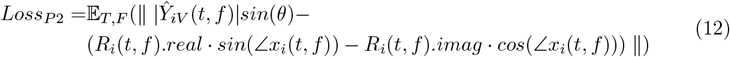

magnitude correlation consistency loss *Loss*_*C*_

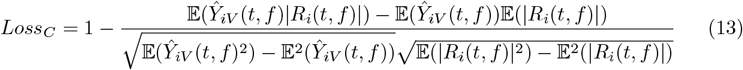

Our full objective is:

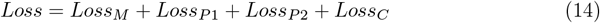

### Overall architecture

The proposed model is shown as Fig 1. It consists of 6 encoder/decoder layers. The first encoder layer consists of 3 directional Inception Resnet blocks. Both the second and third encoder layers consist of an Inception Resnet block and a reduction block. The fourth and fifth encoder layers consist of a reduction block. In each decoder layer, we first used a fractionally-strided convolution with stride 2 and kernel size 2 ×2, batch normalization, LeakyReLU, then used two convolutions with stride 1 and kernel size 3 ×3, batch normalization, LeakyReLU, which followed by 4 Inception-Resnet blocks. In the final layer, we used a 1×1 convolutions and a sigmoid activation function to output 1 channel mask. The mask consists of three submasks: 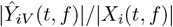, *cos*(*θ*), and *sin*(*θ*).

**Fig 1.**
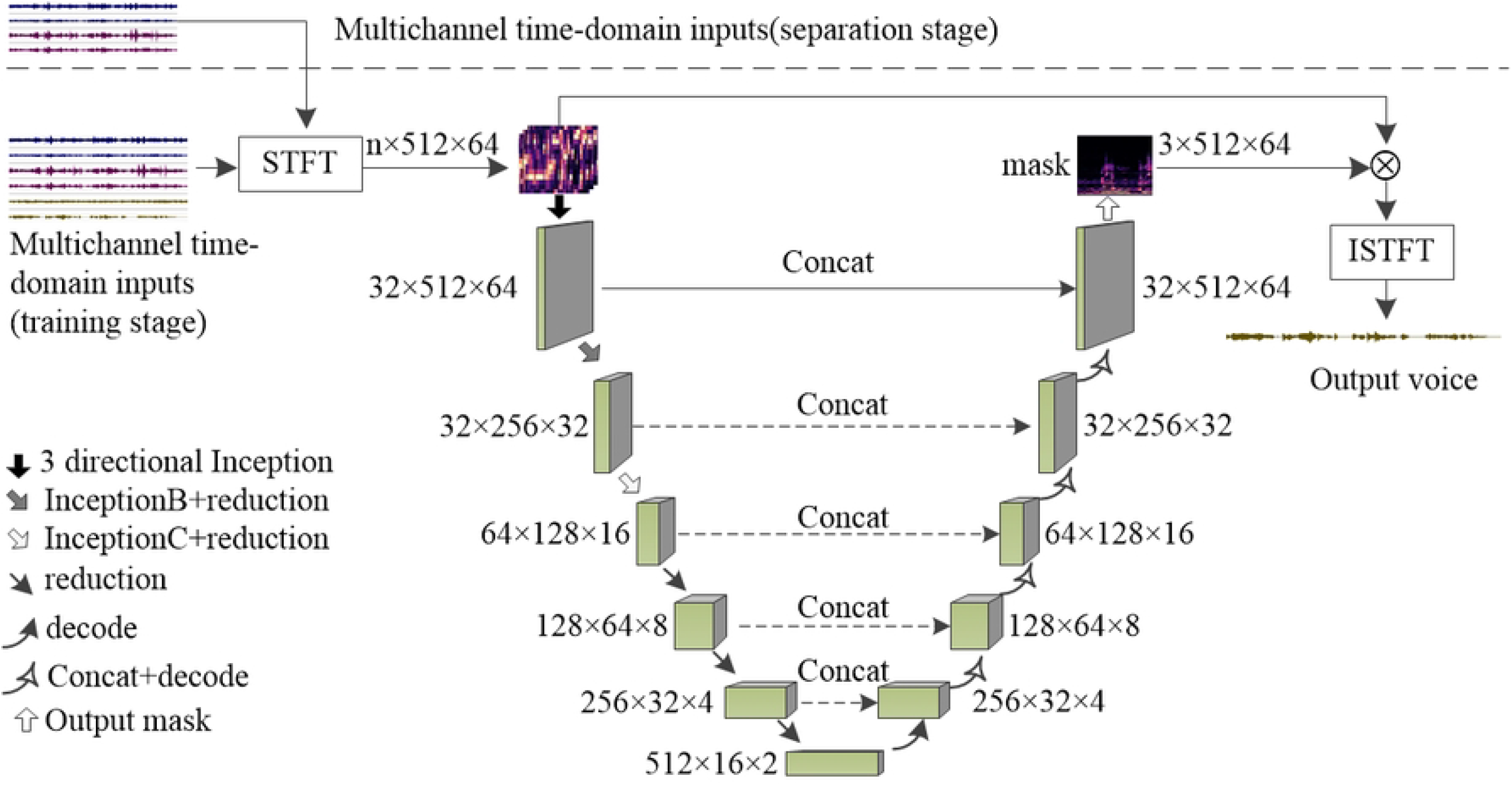
the proposed model

In our approach, songs were recorded at a sample rate of 16,000Hz. We used a 1024-point window size and a 512-point hop size in STFT. Namely, a time window is 64ms, and the overlap between two consecutive windows is 32ms. Thus there are 32 time windows in 1s. Each window is transformed by STFT, generating complex coefficients of 512 valid positive frequency channels between 0 and 8,000Hz. Therefore, the signal lasting 2s will be transformed to a 512 × 64 spectrogram.

## 3 directional Inception-Resnet blocks

The detailed structure of the 3 directional Inception-Resnet blocks is shown in Fig 2. The implementations of the two horizontally oriented Inception-Resnet blocks are similar to that of [36] [37], each 5×5 2D convolution is replaced by two 3×3 convolutions. The flip block in Fig 2 represents the flipping operation of the spectrograms in the horizontal direction. We used a scaling factor of 0.17 to scale the residuals. 10 iterations of the InceptionA block were used in the horizontal direction to cover the spectrograms. The vertically oriented Inception-Resnet block, which is implemented by 3D convolution to extract the spatial features of the multichannel spectrograms, also includes 3 branches: 1×1 convolution, 3×3 convolution, and 5×5 convolution.

**Fig 2.**
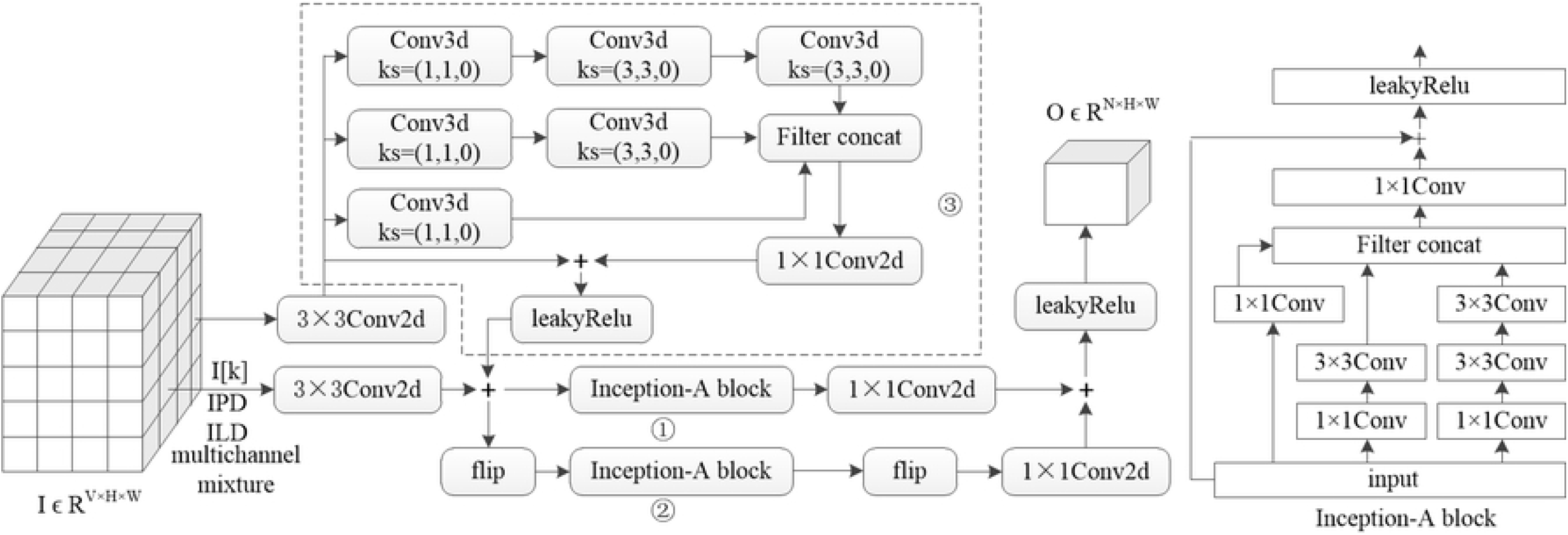
3 directional Inception-Resnet block

### Inception and reduction layers

The Inception Resnet blocks used in the second and third encoder layers are shown in Fig 3. Each block is followed by a reduction filter. To reduce the computational cost, we use a 1×n convolution and a n×1 convolution to replace a n×n convolution in the Inception-B and Inception-C blocks. In the Inception-B and Inception-C blocks, a 1×7 convolution followed by a 7×1 convolution and a 1×3 convolution followed by a 3×1 convolution are used to replace 7×7 convolution and 3×3 convolution, respectively. Each block is iterated 5 times to cover the entire spectrograms. The scaling factor is set to 0.2 in the Inception-B and Inception-C blocks.

**Fig 3.**
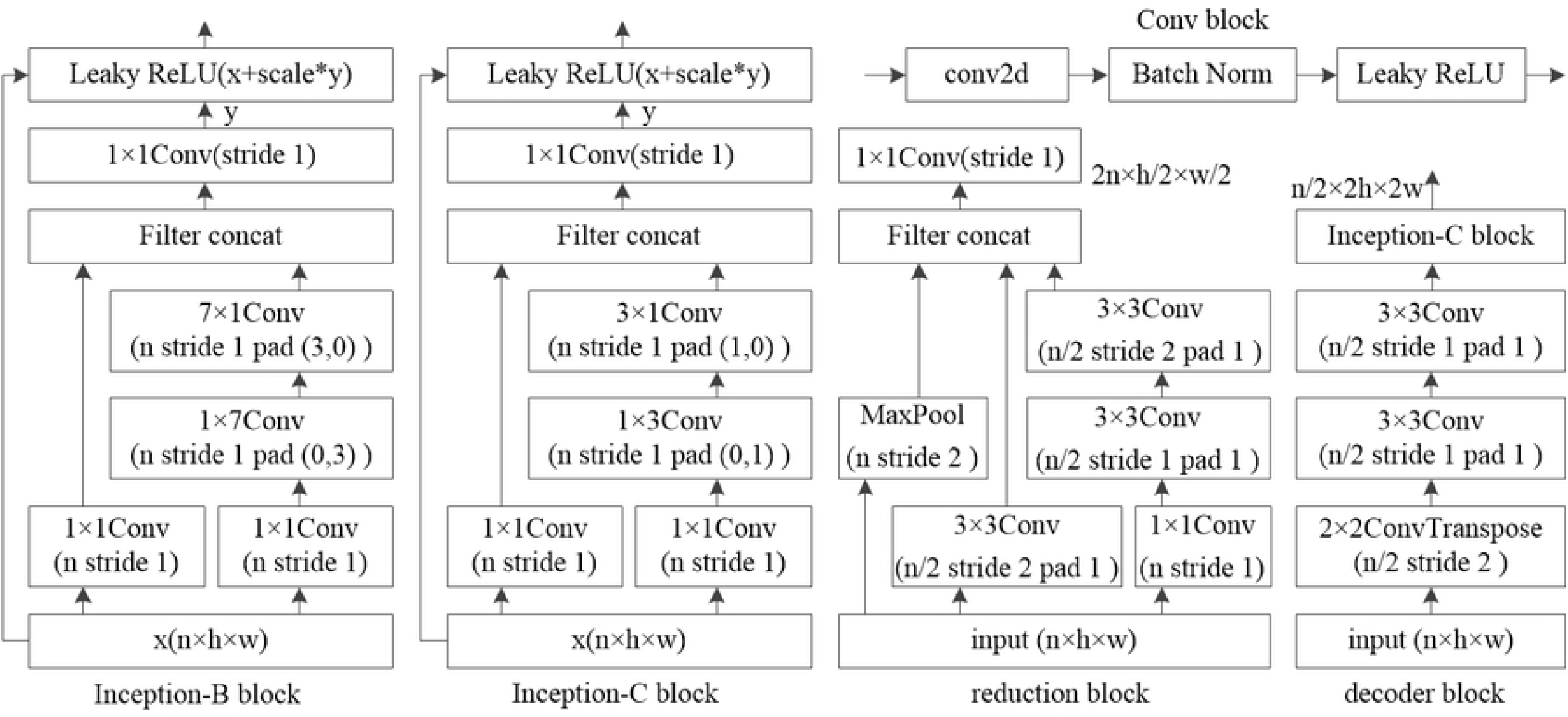
the structure of Inception-B,Inception-C,reduction, and decoder blocks

Convolutional networks usually use maximum or average pooling operations to reduce the size of the activation map. Maximum and average poolings are fast and memory-efficient but lose some information in the activation map. To avoid a representational bottleneck, a series of pooling methods such as power average pooling, stochastic pooling, local Importance Pooling, and softpool have been proposed [38]. Our implementation of the reduction block in each encoder layer is similar to that of [36]. Two parallel 3×3 convolutions with stride 2 are concatenated as in Fig 3. One of the reduction blocks expands the filter banks to avoid the representational bottleneck [37].

### Experiment

#### Parepare Datasets

In our experiment, the 4-channel acoustic signals were recorded by the NAO robot from the public MIR-1K datasets. All song clips in the MIR-1K datasets are sampled at 16000Hz. Each clip is stereo, with one channel for singing voice and another for accompaniment. The 1000 clips were sung by 11 male and 8 female amateurs [31].

The recording scenarios are shown in Fig 4. The recording equipment includes a computer, a robot, and an external speaker. The computer connects the NAO robot via a wireless network. The external speaker that plays the singing voice is placed in front of the robot’s head. The computer plays the singing voice, controls the robot to play the accompaniment, and records the mixture. There are three types of delay during recording: network transmission delay, sound propagation delay, and processing delay. As the singing voice and the accompaniment are played on the computer and the robot respectively, the sound propagation delay of the two sources is different. When recordings, singing voices, and accompaniments are combined into training datasets, they must be aligned.

**Fig 4.**
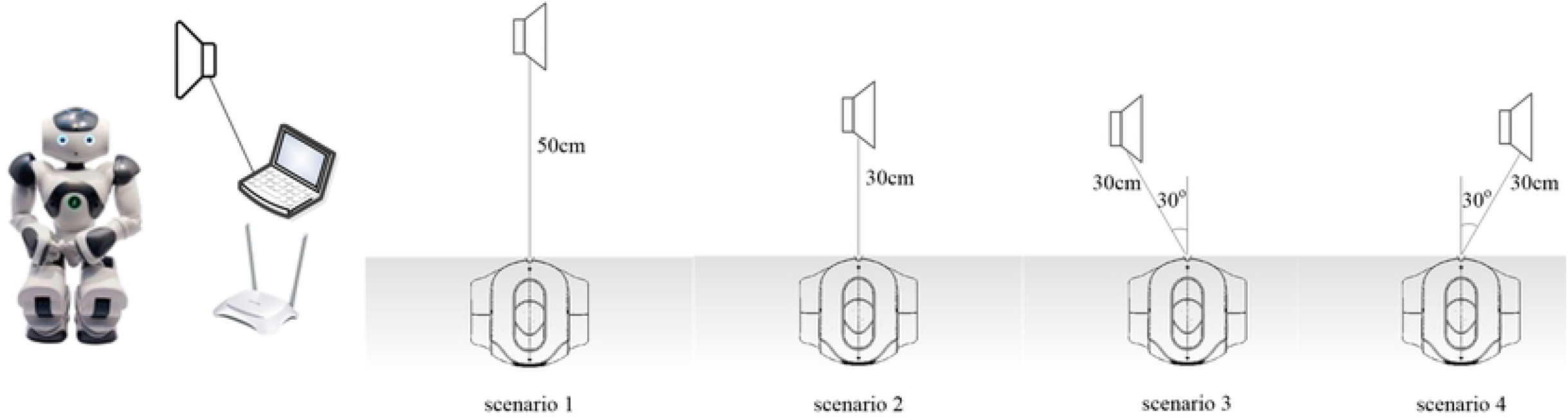
Recording scenarios.

Let *R*_*i*_ denote the *ith* channel recordings, *V* denote the singing voice.

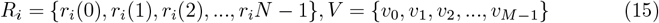

Let 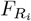 (*p*) denote the recording fragment starting with *p, F*_*V*_ (*q*) denote the singing voice fragment starting with *q*.

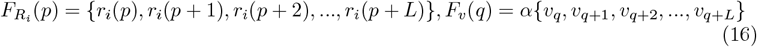

where *L* is the sliding window size and *α* is a coefficient between 0.0-1.0. When 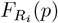 and *F*_*V*_ (*q*) are aligned, *p* and *q* are calculated as follows:

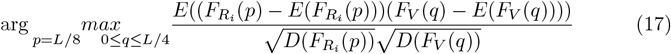

where *E* is the mathematical mean, 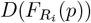 and *D*(*F*_*V*_ (*q*)) are standard deviation of 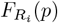 and standard deviation of *F*_*V*_ (*q*).

In our experiment, the song clips were recorded in an unshielded lab. The background noises also included the noise generated by the fan on the robot’s head. The song clips were sampled at 16000 Hz. *L* was set to 64000. As the distances between the microphones on the robot’s head are less than 12 cm, the delay deviations between the microphones are less than 0.3ms or 5 sampling times. The experiments showed that in scenario 1, most delay deviations are 3 sampling times. In this paper, we calculated *p* and *q* of each channel. Clips with a *q* deviation between 2 channels exceeding 6 sampling times were discarded. The training datasets included 2211 aligned 10-channel clips.

The SNR of the singing voice in different scenarios is shown in Fig 5. The mean SNR of the singing voice recorded by the second microphone was the largest. The second microphone was chosen as the reference microphone in our experiment.

**Fig 5.**
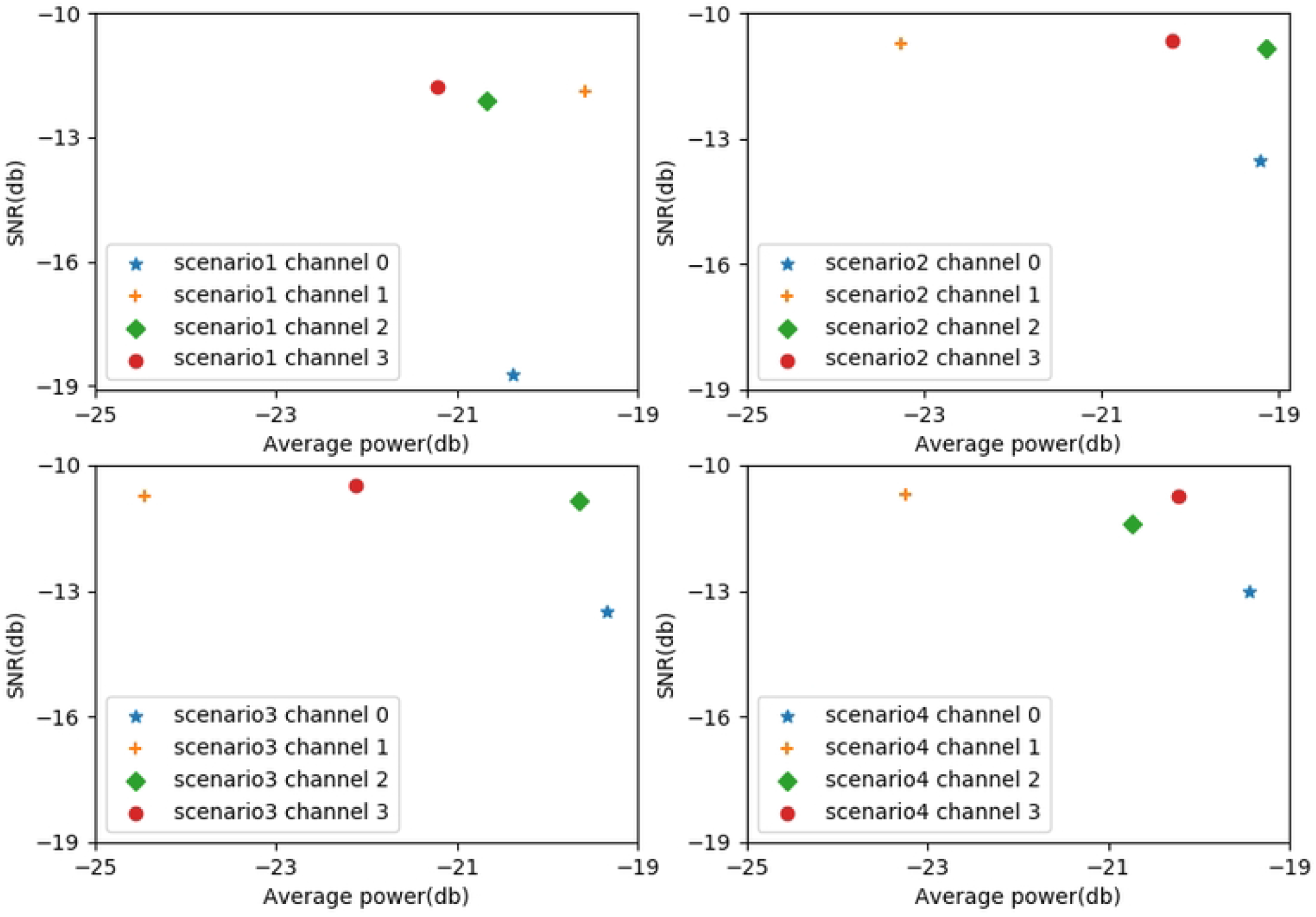
the SNR distribution of the datasets

#### Evaluation

To evaluate the quality of separation, the source to distortion ratio (SDR), source to interferences ratio (SIR), and sources to artifacts ratio (SAR) were taken as objective evaluation criteria [43]. A higher value for each ratio indicates better separation. We used the BSS EVAL toolbox to calculate SDR, SIR, and SAR. We also calculate the normalized SDR (NSDR) provided in the BSS EVAL toolbox.

#### Trainning

In training, the magnitude spectrogram of the mixture was used as the network input, and the magnitude spectrogram of the singing voice was used as the target in the loss function to measure the gap between the predicted result and the singing voice. To evaluate the performance of the proposed model, We trained the proposed model on our datasets. We used 10 Inception-A blocks, 5 Inception-B blocks, and 5 Inception-C blocks in the proposed model. 448 clips were randomly selected in the datasets for training. We randomly selected 448 clips from the 2211 clips for training and 643 clips for testing the performance of the proposed model, and these clips contained each singer’s singing voice in different scenarios. We trained each network for 100 epochs. The optimizer is ADAM. The learning rate is set to 0.00001. The batch size is set to 8. To compare the performance with other models, we also trained the model on the MUSDB and MIR-1K datasets for monaural singing voice separation.

## Results

The performance of the proposed model was evaluated on our datasets, MUSDB datasets, and MIR-1K datasets. Table 1 shows the ablation study on the datasets. The model trained with the mixture, IPD, and ILD for the PSM target achieved better performance than the SMM target. The model with the 3 directional Inception-Resnet blocks has better performance than the others. The performance of the model with 1 directional or 2 directional Inception-Resnet blocks can be improved by adding IPD and ILD to the mixture, while the performance improvement is not obvious for the model with 3 directional Inception-Resnet blocks. In other words, the model with 3 directional Inception-Resnet blocks has the ability to extract the spatial features of the spectrograms.

**Table 1.** Ablation experiment results on the datasets.

| model | target | data | Vocal NSDR | Vocal SIR | Vocal SAR |
| --- | --- | --- | --- | --- | --- |
| 3 directional Inception-Resnet blocks | PSM | mixture+IPD+ILD | 11.5462 | 24.2912 | 10.6632 |
| 3 directional Inception-Resnet blocks | SMM | mixture+IPD+ILD | 11.3162 | 23.4763 | 10.4743 |
| 3 directional Inception-Resnet blocks | SMM | mixture | 11.3171 | 23.4681 | 10.4778 |
| 2 directional Inception-Resnet blocks (①+③) | SMM | mixture+IPD+ILD | 11.1824 | 23.8703 | 10.3046 |
| 2 directional Inception-Resnet blocks(①+②) | SMM | mixture | 11.0004 | 22.4257 | 10.2268 |
| 1 directional Inception-Resnet blocks(①) | SMM | mixture+IPD+ILD | 11.13554 | 22.5646 | 10.3663 |

The separation results for four scenarios are shown in Fig 6. The means of NSDR and SAR in scenario 1 are the lowest. The main reason is that the SNR in scenario 1 is the lowest. The mean NSDR, SIR, and SAR for the other three scenarios were relatively close. The results show that the model can effectively separate the singing voice in different directions.

**Fig 6.**
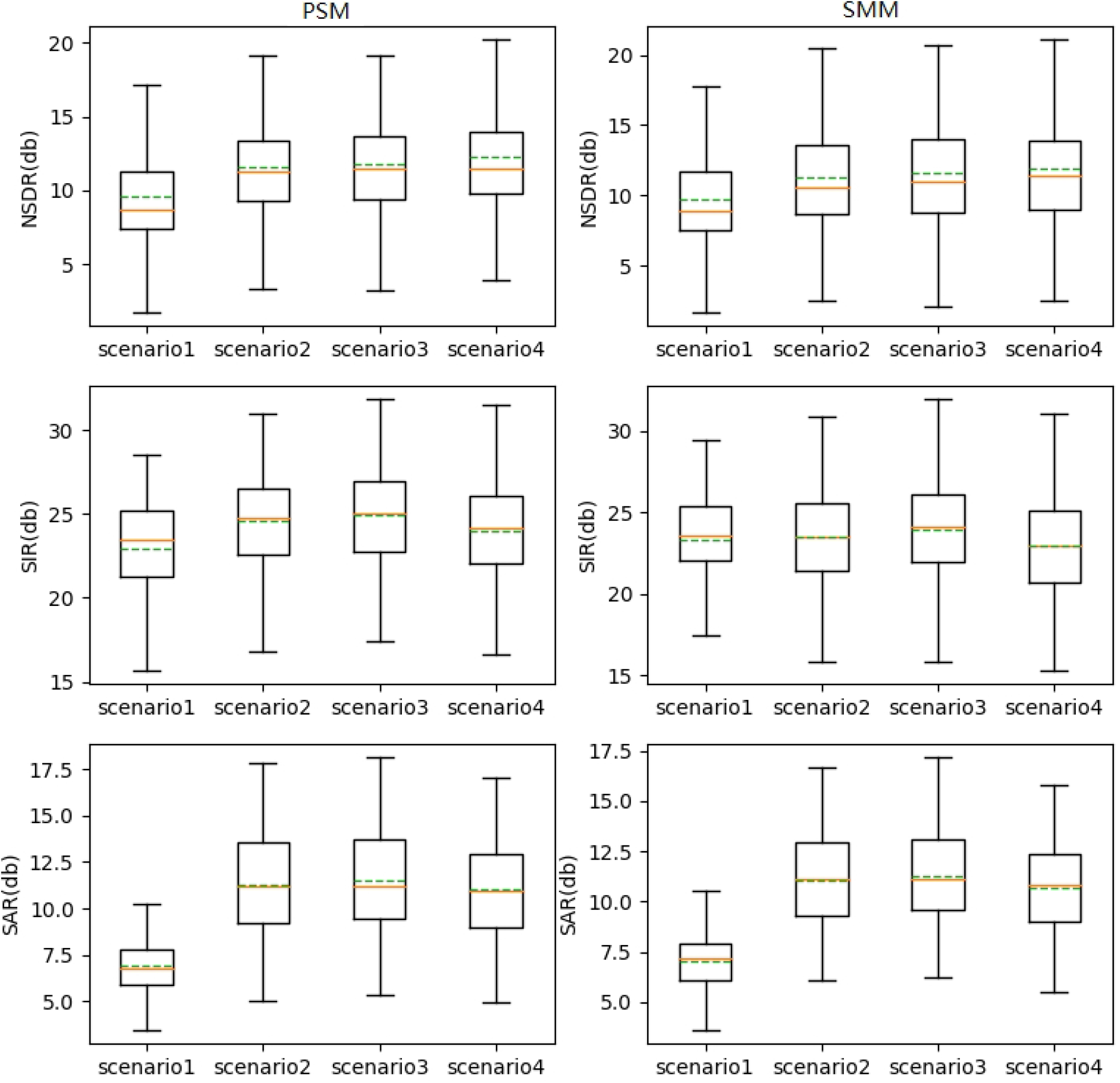
Separation performance of the model

Table 2 shows the separation performance on the datasets with different objectives. We can see that the model trained with *Loss*_*M*_, *Loss*_*P*_, and *Loss*_*C*_ objectives achieved higher NSDR, SIR, and SAR than the two objectives. *Loss*_*C*_ objective can significantly improve the mean NSDR and SAR on the datasets. Table 3 compares the separation performance of our proposed model with the U-Net model. Both models are trained on SMM. The U-Net model was trained with *Loss*_*M*_ objective like [6]. The results show that the proposed model achieved higher NSDR, SIR, and SAR than the U-Net. Due to the distortion of the singing voice recorded by a robot, the U-Net model, which is good at separating the undistorted singing voice, does not achieve a higher separation performance.

**Table 2.** Singing voice mean score on the datasets with different objective.

| objective | Vocal NSDR | Vocal SIR | Vocal SAR |
| --- | --- | --- | --- |
| $Loss_M + Loss_P + Loss_C$ | 11.3171 | 23.4681 | 10.4778 |
| $Loss_M + Loss_P$ [48] | 9.3305 | 23.2536 | 8.4747 |

**Table 3.** Singing voice mean score on the datasets for different model.

| model | Vocal NSDR | Vocal SIR | Vocal SAR |
| --- | --- | --- | --- |
| Inception-ResUnet | 11.3171 | 23.4681 | 10.4778 |
| U-Net [6] | 6.5867 | 17.4178 | 5.9888 |

To compare the performance of Inception-ResUNet with other models, we also trained the Inception-ResUNet model on MUSDB datasets and MIR-1K datasets with 2 directional Inception-Resnet blocks, PSM, and *Loss*_*M*_ + *Loss*_*P*_ . Table 4 shows the comparison of Inception-ResUNet with other models for SDR, SIR, and SAR on MUSDB datasets.

**Table 4.**
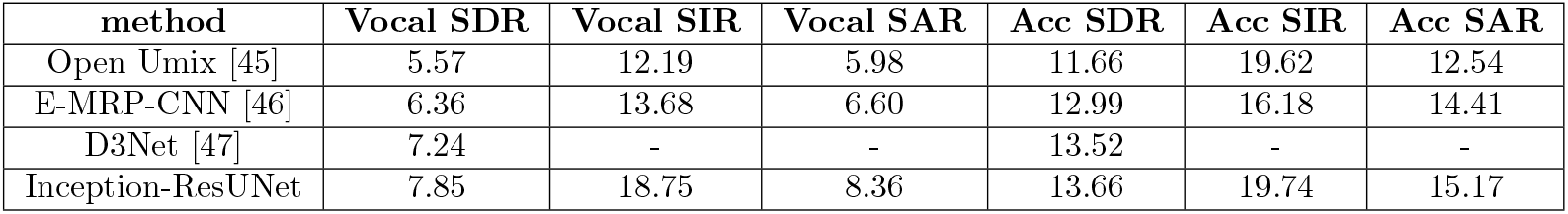
Comparison of SDR,SIR,SAR of other methods and Inception-ResUnet on MUSDB datasets.

As shown in Table 4, the proposed model achieves 7.85db on the vocal SDR category and 13.66db on the accompaniment SDR category on the MUSDB datasets, which outperforms Open Umix, E-MRP-CNN, and D3Net. As shown in Table 5, the proposed model achieves 12.73db on the vocal NSDR category and 12.53db on the accompaniment NSDR category on the MIR-1K datasets, which outperforms E-MRP-CNN, U-Net, and RPCA-DRNN.

**Table 5.**
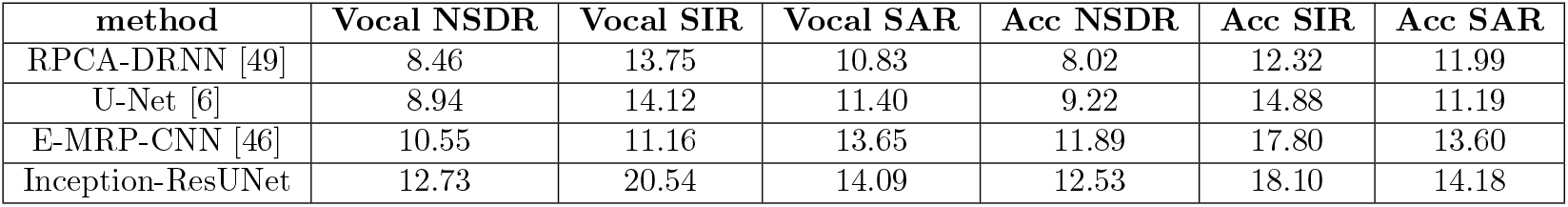
Comparison of NSDR,SIR,SAR of other methods and Inception-ResUnet on MIR-1K datasets.

## Discussion

The separation performances of most singing voice separation methods for monaural recordings degrade for separating distorted singing voices. The main reason is that the distortion of the spectrogram is also proportionally reserved. Unfortunately, the multichannel mixture recorded by ordinary robots is distorted, as shown in position 1 in Fig 7. The model trained by *Loss*_*M*_ and *Loss*_*P*_ [6] [46] [48] will preserve the distortion, as shown in position 2 in Fig 7. When a model is trained on multi-objectives, improvements of one objective will degrade the others. The proposed model trained by *Loss*_*M*_ + *Loss*_*P*_ + *loss*_*C*_ can not achieve the best separation performance on MUSDB18 datasets. Our experiments show that a model trained by *Loss*_*M*_ + *Loss*_*P*_ + *loss*_*C*_ had a lower SDR. However, when *Loss*_*C*_ is used to separate distorted singing voices, *Loss*_*C*_ can significantly improve separation performance. *loss*_*C*_ reduces the distortion of the singing voice, and the benefits outweigh the reduction.

**Fig 7.**
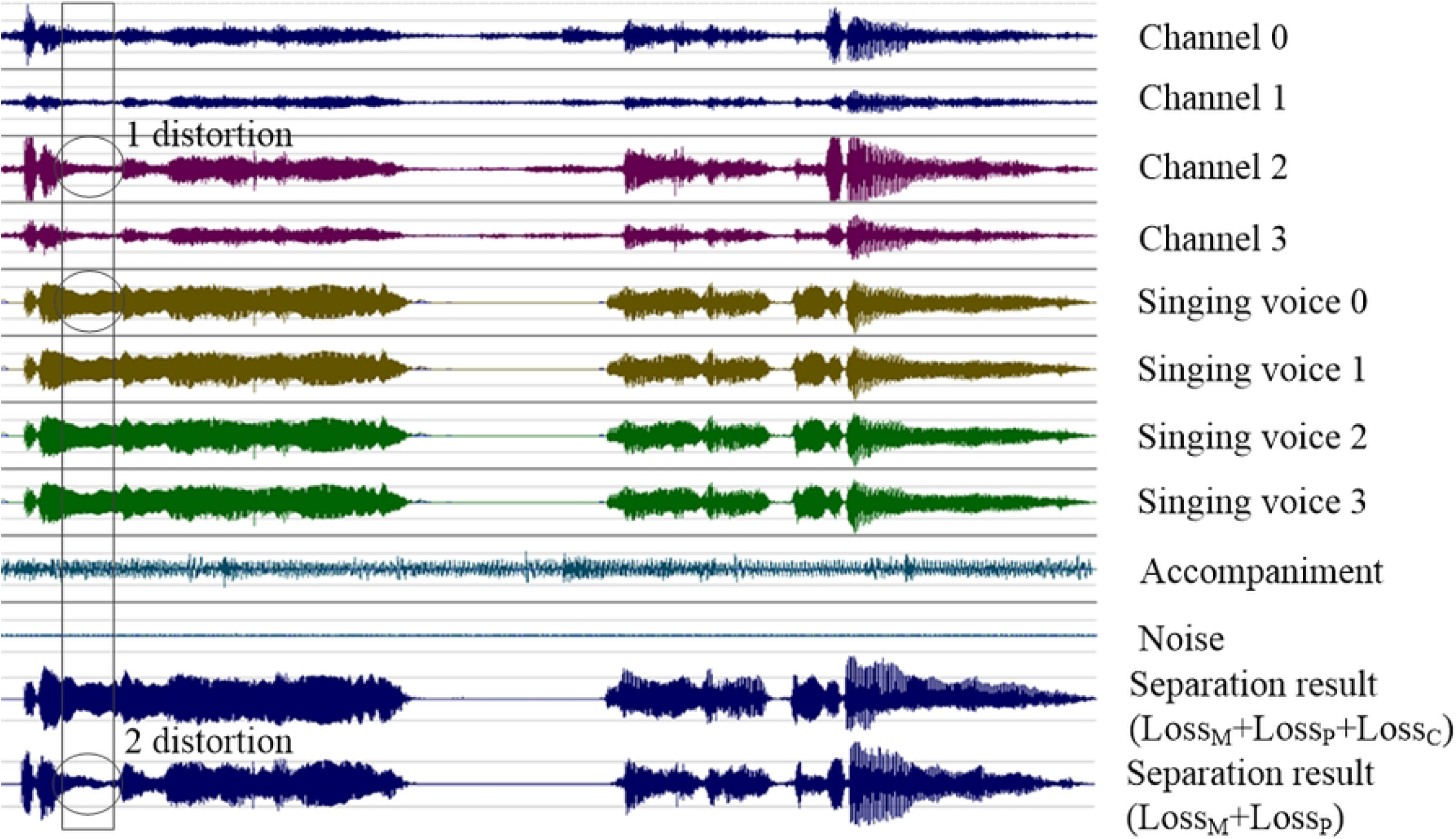
*Loss*_*C*_ corrects the distortion of singing voice separation

## Conclusion

In this paper, We proposed a novel model, 3 directional Inception-ResUNet, for separating the multichannel singing voice with distortion. We trained the proposed model with multi-objectives: magnitude correlation consistency loss, magnitude consistency loss, and phase consistency loss. We recorded multichannel singing voices on robots and produced 10-channel datasets to test multichannel singing voice separation algorithms. The output of the proposed model is a set of masks of the singing voice that can be used to transform the magnitude and phase spectrograms of the mixture into the singing voice. The experimental results show that the proposed model achieved higher performance on the multichannel singing voice separation with distortion.

